# Characterization of SOX2, OCT4 and NANOG in ovarian cancer tumor-initiating cells

**DOI:** 10.1101/2020.09.08.288381

**Authors:** Mikella Robinson, Samuel F Gilbert, Jennifer A Waters, Omar Lujano-Olazaba, Jacqueline Lara, Logan J Alexander, Samuel E Green, Gregory Burkeen, Omid Patrus, Ryne Holmberg, Christine Wang, Carrie D House

## Abstract

Identification of tumor initiating cells (TICs) has traditionally relied on expression of surface markers such as CD133, CD44, and CD117 and enzymes such as aldehyde dehydrogenase (ALDH). Unfortunately, these markers are often cell type specific and not reproducible across patient samples. A more reliable indication of TICs may include elevated expression of stem cell transcription factors such as SOX2, OCT4, and NANOG that function to support long-term self-renewal, multipotency, and quiescence. RNA-sequencing studies presented here highlight a potential role for SOX2 in cell cycle progression in cells grown as 3-D spheroids, which are more tumorigenic and contain higher numbers of TICs than their 2-D monolayer cultured counterparts. SOX2, OCT4, and NANOG have not been comprehensively evaluated in ovarian cancer cell lines, although their expression is often associated with tumorigenic cells. We hypothesize that SOX2, OCT4, and NANOG will be enriched in ovarian TICs and will correlate with chemotherapy resistance, tumor initiation, and expression of traditional TIC markers. To investigate this hypothesis, we evaluated SOX2, OCT4, and NANOG in a panel of eight ovarian cancer cell lines grown as a monolayer in standard 2-D culture or as spheroids in TIC-enriching 3-D culture. Our data show that the high-grade serous ovarian cancer (HGSOC) lines CAOV3, CAOV4, OVCAR4, and OVCAR8 had longer doubling-times, greater resistance to chemotherapies, and significantly increased expression of SOX2, OCT4, and NANOG in TIC-enriching 3-D culture conditions. We also found that in vitro chemotherapy treatment enriches for cells with significantly higher expression of SOX2. We further show that the traditional TIC marker, CD117 identifies ovarian cancer cells with enhanced SOX2, OCT4, and NANOG expression. Tumor-initiation studies and analysis of The Cancer Genome Atlas (TCGA) suggest a stronger role for SOX2 in ovarian cancer relapse compared with OCT4 or NANOG. Overall, our study clarifies the expression of SOX2, OCT4, and NANOG in TICs from a variety of ovarian cancer cell lines. Our findings suggest that SOX2 expression is a stronger indicator of ovarian TICs with enhanced tumor-initiation capacity and potential for relapse. Improved identification of ovarian TICs will advance our understanding of TIC biology and facilitate the design of better therapies to eliminate TICs and overcome chemotherapy resistance and disease relapse.

## Introduction

High Grade Serous Ovarian Cancer (HGSOC) is the most lethal gynecological malignancy in women. Although a majority of patients respond to platinum-based chemotherapies, over 70% relapse with therapy-resistant disease within 18 months [1, 2]. Mechanisms underlying ovarian cancer recurrence are unclear but likely involve a small population of relatively undifferentiated tumor-initiating cells (TICs) that can resist chemotherapy and efficiently reestablish heterogeneous tumors. TICs share properties of tissue stem cells such as quiescence, long-term self-renewal, asymmetric division, and differentiation potential [3–7]. Consequently, TICs are slow cycling cells with blocked apoptosis, high drug efflux, and enhanced activation of developmental signaling pathways such as WNT, NOTCH, and Hedgehog [8–13]. Traditional chemotherapeutic drugs, while successful at killing the bulk cells of the tumor, are inefficient at eliminating TICs. Thus, TICs are increasingly recognized as drivers of recurrent disease in many tumor types, with studies focused on identifying and eliminating these elusive cells being crucial for overcoming chemotherapy resistance and preventing relapse [14].

Identification of ovarian TICs has traditionally relied on a combination of cell surface proteins, mainly CD133, CD44, and CD117 as well as the activity of aldehyde dehydrogenase (ALDH) enzymes, including ALDH1A1 and ALDH1A2 [15–18]. Given their role in maintaining pluripotency and long-term self-renewal, it is not surprising that the embryonic transcription factors SOX2, OCT4, and NANOG are recognized as part of the stem cell signature and often correlate with surface and enzyme markers [19, 20]. Indeed, enhanced expression of SOX2, OCT4, and/or NANOG is typical of ovarian TICs, and their expression enhances spheroid formation, drug efflux, chemoresistance, and induction of epithelial to mesenchymal (EMT)-related genes [15, 17, 21–24].

As research has progressed in this field it has become more apparent that TIC populations are heterogeneous, expressing different combinations of markers that are often cell-line or patient specific. For example, we previously found the expression levels of SOX2, OCT4, and NANOG were significantly different between two ovarian cancer cell lines, even though both lines showed similar enhancement of ALDH activity [25]. Bareiss et al., showed that SOX2 gene expression levels varied across five ovarian cancer cell lines and four patient samples [19] and Silva, et al. similarly found variable expression of ALDH, CD133, and CD117 across cell lines and patient samples [16]. The lack of reliable markers is likely due to the genetic heterogeneity of ovarian cancer as well as the stage of disease and tissue of origin for different subsets of ovarian TICs [14, 26–28]. Further complicating the findings are recent genomic analyses demonstrating that some of the most commonly used ovarian cancer cell lines do not exhibit the genetic profile corresponding to the tumor type they are being used to study [29–31]. These findings suggest some cell lines may be unreliable for investigating mechanisms of drug resistance and relapse of high-grade serous ovarian cancer (HGSOC), the most common histological subtype responsible for the majority of fatalities from all ovarian cancers combined [32]. Moreover, there has been no comprehensive study that compares the expression of SOX2, OCT4, and NANOG in the most commonly used ovarian cancer cell lines and whether their expression correlates with specific surface markers and other features of TICs.

To begin to address these deficiencies we investigated the expression of SOX2, OCT4, and NANOG in a panel of ovarian cancer cell lines to identify correlations with growth properties, chemoresistance, tumor-initiating potential, and expression of traditional TIC markers. Our analysis includes five high-grade serous cell lines, two undefined serous lines, and one non-serous line which were cultured as spheroids in 3-D conditions compared relative to standard culture as a monolayer in 2-D conditions. We show that SOX2, OCT4 and NANOG are consistently enriched in HGS lines cultured in 3-D relative to 2-D and their expression correlates with genes encoding traditional TIC markers. Cells expressing CD117 or CD133 in combination with high ALDH activity identifies cells with relatively high SOX2, OCT4, and NANOG expression. Cells with high tumor initiation capacity and resistance to chemotherapy have significantly enriched SOX2, but not OCT4 or NANOG. To our knowledge these data provide the first comprehensive evaluation of SOX2, OCT4 and NANOG in TICs from commonly used ovarian cancer cell lines. Clarification of these important factors will lead to more reliable identification of ovarian TICs and better characterization of pathways regulating drug resistance and other stem-like features.

## Materials and Methods

### Cell Lines and Culture Conditions

OVCAR8 (HTB-161), CAOV3 (HTB-75), CAOV4 (HTB-76), SKOV3 (HTB-77), and OV90 (CRL-11732) were obtained from American Type Culture Collection (ATCC) and maintained in Roswell Park Memorial Institute (RPMI) medium. OVCAR4 and OVCAR5 were obtained from NCI-Frederick DCTD tumor/cell line repository and also maintained in RPMI. ACI23 cell line was provided by Dr. John Risinger and Memorial Health University Medical Center, Inc. and maintained in Dulbecco’s Modified Eagle Nutrient Mixture F-12 (DMEM/F12) medium (Gibco).

All 2-D monolayer cultures (2-D) were maintained in standard RPMI media supplemented with 10% fetal bovine serum (FBS) and 1% penicillin-streptomycin (Gibco) in tissue-culture treated flasks at 37°C with 5% CO2. Except where noted, TIC-enriching 3-D cultures (3-D) were maintained in stem cell media: DMEM:F12 (+ L-glutamine, + 15 mM HEPES) (Gibco) supplemented with 1% penicillin/streptomycin, 1% KnockOut serum replacement (Gibco), 0.4% bovine serum albumin (Sigma-Aldrich), and 0.1% insulin-transferrin-selenium (Gibco) and cultured in non-treated flasks at 37°C with 5% CO2. Cultures using stem cell media were supplemented with human recombinant epidermal growth factor (EGF) and basic fibroblast growth factor (FGF) every 2–3 days for a final concentration of 20 ng/ml and 10 ng/ml, respectively.

### Spheroid formation assay

To generate spheroids, 500 cells/well in 100 μL were seeded in ultra-low attachment flat bottom 96-well plates (Corning) with either standard RPMI media or TIC-culture media. After 7 days in culture, whole wells were stained with Hoechst 33342 (ThermoFisher Scientific) and imaged at 10x magnification using the ImageXpress Pico (Molecular Devices). Spheroids were analyzed using the CellReporterXpress software (Version 2.1.5156, Molecular Devices), selecting for spheroids >30 μm. Spheroid formation efficiency was calculated for cells grown in standard or TIC culture media.

### RNA Extraction and Quantitative Reverse Transcription PCR (qRT-PCR)

Total RNA was isolated using either NucleoSpin RNA Plus purification kit (Macherey-Nagel) or, in the case of FACS-sorted cells, Direct-zol RNA Miniprep Plus kit (Zymo Research) per manufacturer’s instructions. Final RNA concentration was determined using the 260/280 absorbance ratio with a SpectraMax QuickDrop spectrophotometer (Molecular Devices). Total purified RNA was converted to cDNA using the High-Capacity cDNA Reverse Transcription Kit (ThermoFisher Scientific). Analysis of gene expression was performed with Taqman Fast Advanced Master Mix and Taqman probe assays with GAPDH as control (ThermoFisher Scientific [Oct4 Assay ID: Hs04260367_gH, Sox2 Assay ID: Hs01053049_s1, Nanog Assay ID: Hs02387400_g1, GAPDH Assay ID: Hs99999905_m1, CD44 Assay ID: Hs01075862_m1, CD117 Assay ID: Hs00174029_m1, CD133 Assay ID: Hs01009259_m1, ALDH1A1 Assay ID: Hs00946916_m1, ALDH1A2 Assay ID: Hs00180254_m1). Reactions were run on the QuantStudio 3 and analyzed on the QuantStudio 3 Design and Analysis software v1.5.1 (ThermoFisher Scientific). Quantitation and normalization of relative gene expression were accomplished using comparative threshold cycle method or ΔΔCT.

### Cell viability Assay

Cell viability was assessed with CellTiter-Glo luminescent reagent (Promega) according to manufacturer’s instructions using a white, 96-well plate. SpectraMax iD3 plate reader and SoftMax Pro Software 7.1.0 (Molecular Devices) was used to quantify cell viability relative to control wells.

### Flow Cytometry

Cells were grown for 5 days in TIC-enriching conditions. Spheroids were prepared into single cell suspensions using Cellstripper (Corning) and needle dissociation. Fluorescently-conjugated antibodies were purchased from Cell Signaling Technology: CD44-FITC (Cat. no. 3570S) or Miltenyi Biotec: CD117-APC (Cat. no. 130-111-671), CD133-FITC (Cat. no. 130-110-968). CD44 was used at 1:10000 and CD117 and CD133 at 1:100 dilution for 1 hr at 4°C. ALDH activity was assessed using the AldeRed Kit (Millipore Sigma – Cat. No. scr150) according to manufacturer’s instructions. Sorting of positive and negative cells was performed on a BDFACS Melody cell sorter (Becton Dickinson) directly into RNA lysis buffer and immediately processed for qRT-PCR studies. Gates provided in **Supplemental Figure 1**. Cell percentages were analyzed using BD FACSChorus Software (Becton Dickinson).

### Animal Experiments

All animal studies were approved by the SDSU Animal Care and Use Committee. For subcutaneous xenografts, 500,000 to 500 ACI23 cells in 200 ul of 1:1 Matrigel and PBS were subcutaneously injected into the left flank of 8-week-old female athymic Nu/Nu mice. For controls, 200 ul of 1:1 Matrigel and PBS was injected into the right flank of each mouse. Mice were weighed and tumors measured using calipers twice weekly. Mice were monitored for 120 days and sacrificed once tumors reached 20mm length or 120 days. Only mice that developed tumors were compared. Early and Late groups were created for each dilution by ranking tumors based on how many days it took for palpable tumors to appear (time to tumor initiation) and splitting into two equal groups.

### Immunohistochemistry (IHC) and Immunocytochemistry (ICC)

For IHC, tumors were fixed at excision in 10% neutral buffered formalin and stored in 70% ethanol before processing. Tumors were embedded in paraffin and sectioned at 5um. Antigen retrieval was performed in the presence of citrate buffer (pH 6). Slides were quenched (3% hydrogen peroxide, 5 min) and blocked (5% normal goat serum, 30 min). Slides were incubated overnight at 4°C with primary antibodies purchased from Cell Signaling Technology: SOX2 1:300 (Cat. no. 3579S), OCT4 1:200 (Cat. no. 2750S), and NANOG 1:600 (Cat no. 4903S) followed by an HRP-linked polymer (Cat. no. 8114S) for 1 hour at room temperature. Slides were processed using the DAB (3,3’-Diaminobenzidine) kit (Vector) for 10 minutes. Four randomly selected images per slide were acquired with a Carl Zeiss Primo Star HAL/LED Microscope and imaged using Toupview imaging program. Three blinded investigators scored the immunostainings based on the staining intensity in the positively stained areas, as described [33].

### Public Database Analysis

The prognostic significance of *SOX2, OCT4* (*POU5F1*), and *NANOG* genes was determined by comparing their mRNA expression and Ovarian Serous Cystadenocarcinoma disease recurrence using The Cancer Genome Atlas (TCGA) data provided in the Cancer Genomics Portal Website (www.cbioportal.org) [34, 35]. Each gene was analyzed for gene expression z-score, associated with survival outcomes, and reconstructed in an excel spreadsheet by unique patient ID. There were 579 total cases, where 295 were complete with mRNA expression z-scores (relative to all cells “log RNA Seq V2 RSEM”) and 137 were also coded for recurrent dead, recurrent alive, and disease free alive.

### Statistical Analysis

Statistics were generated using Prism7 (GraphPad) with data acquired from at least three independent biological replicates. Results are presented as mean +/- SEM. Significance was calculated using either student’s t-test for comparisons of two means or ANOVA for comparisons of three or more means with a Tukey post hoc test to identify differences between groups or as described in figure legends. Correlation comparisons were analyzed using Pearson’s Correlation. Differences between means are considered statistically significant at the 95% level (P<0.05).

## Results

In this study we sought to characterize TIC markers and corresponding stem cell transcription factors in a panel of ovarian cancer cell lines with different growth properties. We evaluated commonly used cell lines that were defined by genetic analysis as possibly or likely HGSOC (OV90/CAOV3/CAOV4 /OVCAR4/OVCAR8) or unlikely HGSOC (SKOV3) [30]. We also included two undefined serous ovarian cancer lines (ACI23, OVCAR5) [36, 37].

We previously showed that ovarian cancer cells cultured in 3-D conditions enhances growth of tumor cells as multicellular spheroids and enriches for TICs with elevated expression of stem cell markers, increased tumor-initiating capacity, and greater resistance to chemotherapies [15, 25]. Subsequent RNA-sequencing and Gene Set Enrichment Analysis (GSEA) showed an increase in genes involved in drug metabolism and oxidative stress in OV90 cells cultured in 3-D relative to 2-D conditions [38]. A further query of this sequencing data identifies 10,222 significantly differentially expressed genes (DEGs) in the 3-D cultured cells relative to 2-D cultured cells (**Fig. 1A**). DEGs representing a log fold change of at least two, of which there were 4,045 genes, are indicated in red in the volcano plot. It is noteworthy that included in these DEGs were *SOX2*, and *ALDH1A2* which are significantly increased in 3-D conditions relative to 2-D. DEGs representing a log fold change of at least one but less than two are indicated in blue and include the stem cell markers *CD44* and *CD133* (*Prom1*) in 3-D conditions relative to 2-D (**Fig. 1A**). There was no significant difference in expression of *OCT4, NANOG, CD117*, or *ALDH1A1*.

**Figure 1.**
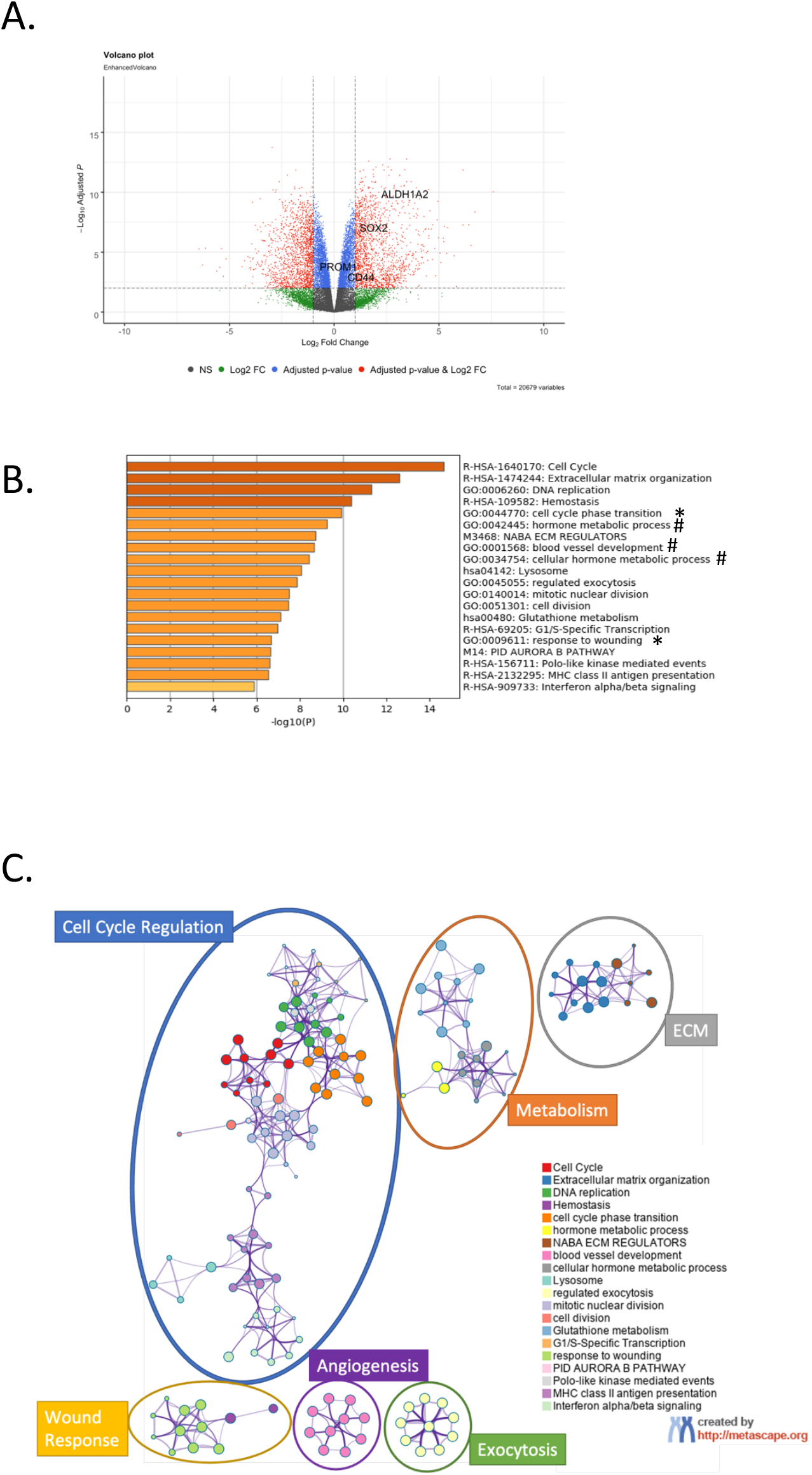
RNA-Sequencing of ovarian cancer cells cultured in 3-D conditions relative to 2-D cultures. A) Volcano plot of RNAseq DEGs. Blue genes adj p<0.05, Red genes Log2FC<1 and adj p<0.05; B) Metascape GO analysis of DEGs Log2FC<-1.2 or Log2FC>1.2 and adj p<0.000001, terms labelled * include SOX2, # include ALDH1A2; C) Metascape GO tree showing GO clusters.

To identify pathways that support cell growth in 3-D conditions, the top 10% DEGs were selected for gene ontology (GO) studies. DEGs with an adjusted P value < 0.000001 and log2FC>1.2 or log2FC<-1.2 were subjected to Metascape gene annotation and analysis. Metascape analysis indicated several cell cycle regulation pathways are altered in 3-D conditions, including cell cycle, extracellular matrix organization, DNA replication, cell cycle phase transition, and wound healing, among several others (**Fig. 1B**). *SOX2* gene (noted by *) appears in cell cycle phase transition and wound healing, while *ALDH1A2* gene (noted by #) appears in blood vessel development and metabolic process pathways. A Metascape GO tree shows that the cell cycle regulation terms cluster together while the metabolism regulation terms also cluster together (**Fig. 1C**). These data suggest that in addition to altered metabolism and oxidative stress, which we have previously shown to support ovarian cancer spheroids [38], cell cycle regulation pathways play a critical role in supporting growth of ovarian cancer cells in 3-D and may correlate with specific markers of TICs.

Given the heterogeneity of ovarian cancer cell lines [18, 39–41], we first evaluated growth properties of a panel of commonly reported lines cultured in 2-D and 3-D conditions. Standard 2-D culture conditions revealed differential growth over a 7-day period among the cell lines (**Fig. 2A**). Growth was slower in 3-D conditions for all cell lines except ACI23, OVCAR5, and OV90, which had increased growth relative to 2-D conditions (**Fig. 2B-C**). ACI23 and OVCAR8 had the shortest doubling time of ~1.66 days each, whereas OVCAR4 had the longest doubling time of ~3.54 days in 2-D culture (**Fig. 2C**). In accordance with their growth in 2-D, ACI23 cells had the shortest doubling time and OVCAR4 cells had the longest doubling time in 3-D culture (**Fig. 2C**). The shorter doubling of ACI23, OVCAR5, and OV90 cells in 3-D relative to 2-D suggest these lines have less dependence on serum and anchorage support for growth.

**Figure 2.**
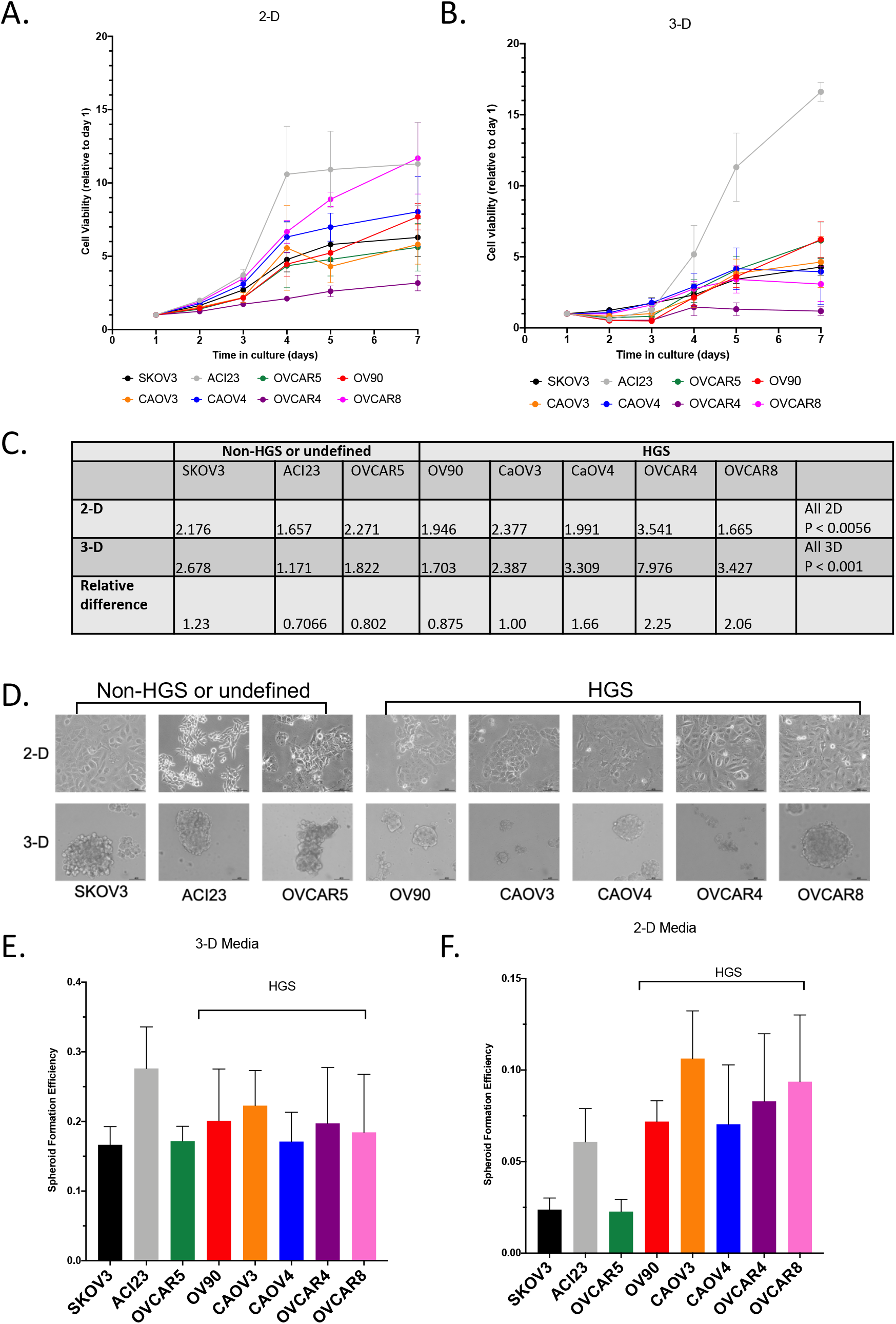
Growth properties of ovarian cancer cells cultured in 2-D or 3-D conditions. Viability growth assay of cells grown over 7 days for A) 2-D conditions and B) 3-D conditions; C) Doubling time for 2-D and 3-D conditions, calculated with Least Squares Fit of Log Exponential Growth. Sum of Squares Test; D) Bright Field Images of OVCA cells in 2-D and 3-D conditions, 10x; Spheroid Formation Efficiency for OVCA cells grown on ultra-low attachment plates in E) 3-D media and F) 2-D media. Graphs represent mean and SEM.

We next measured spheroid formation efficiency in TIC enriching 3-D conditions. Spheroids are multicellular tumor cell aggregates that mimic those found in patient ascites and exhibit enhanced drug resistance and high tumorigenicity [15, 38, 42, 43]. Spheroid formation is an in vitro surrogate used to identify cell lines with potentially high tumor-initiation capacity [44–46]. All cell lines had relatively equal ability to form spheroids in 3-D conditions (**Fig. 2D-E**). The high sphere-forming efficiency of ACI23 cells may be attributed to its ability to thrive under serum deprived, low anchorage culture conditions (**Fig. 2A**). Given the varying response of ovarian cancer cells to growth factor stimulation [41], we also compared spheroid formation ability in ultra-low attachment plates using standard 2D culture media, which contains serum and lacks EGF and FGF. We found that all cell lines exhibited lower spheroid formation efficiency when cultured in standard 2-D media (**Fig. 2E**). The average efficiency across all cell lines in 3-D media was ~0.19 whereas this was reduced to ~0.067 in standard 2-D media. The SKOV3 and OVCAR5 cell lines appear to be most dependent on EGF and FGF as they had the lowest efficiency with standard 2-D media, although neither was statistically significantly different compared to the other cell lines (**Fig. 2F**). Interestingly, cell lines defined as HGS all exhibit comparatively similar spheroid formation efficiency whether in 2-D or 3-D.

In order to better clarify the involvement of SOX2, OCT4 and NANOG in 3-D growth of ovarian cancer cells and to validate the RNA-sequencing data, we evaluated the expression levels of these genes in 3-D relative to 2-D conditions in our panel of cell lines. Although all three embryonic transcription factors are known to support pluripotency and long-term self-renewal, to our knowledge there has been no comprehensive analysis of SOX2, OCT4, and NANOG levels in 3-D cultures or in TICs sorted from commonly used ovarian cancer lines. All cell lines had increased expression of SOX2, OCT4 and NANOG when cultured in 3-D relative to 2-D conditions although the HGS lines CAOV3, CAOV4, OVCAR4, and OVCAR8 and the undefined ACI23 line had the greatest enrichment of all three genes (**Fig. 3A-C**). Although OV90 is likely HGS [29, 30], 3-D culture of this line did not enhance expression of SOX2, OCT4, and NANOG to the same levels as the other HGS lines included in this study (**Fig. 3A-C**). SKOV3 and OVCAR5 cells had no statistically significant increase in expression of either gene, suggesting they are less dependent on SOX2, OCT4, and NANOG for growth in 3-D. We next quantified the transcript levels of CD44, CD133, CD117, ALDH1A1 and ALDH1A2, traditionally used markers of ovarian TICs, in 3-D relative to 2-D cultures in our cell line panel. Our data show that, in contrast to SOX2, OCT4, and NANOG, ovarian TIC marker gene expression is more variable across cell lines (**Fig. 3D-H**). Although most lines exhibited increased levels of CD44, CD133, ALDH1A1 and ALDH1A2 in 3-D relative to 2-D (**Fig. 3D and 3F-G**), the opposite was true for CD117 (**Fig. 3E**). With the exception of CAOV4, all cell lines showed no change or decreased expression of CD117 in 3-D relative to 2-D. Moreover, unlike ALDH1A1 which was not significantly enriched in any cell line cultured in 3-D, ALDH1A2 was significantly higher in ACI23, OV90, and CAOV3 cells cultured in 3-D. The difference in ALDH1A1 and ALDH1A2 expression across cell lines may indicate cell-line specific dependence on distinct ALDH isoforms. A correlation analysis revealed that CD44, CD117, and CD133 significantly correlate with expression of SOX2, OCT4, and NANOG in 3-D relative to 2-D cultures (**Fig. 3I**). ALDH1A1 significantly correlates with NANOG expression whereas ALDH1A2 does not correlate with any other genes in this analysis. Taken together these data highlight the heterogeneous expression of TIC marker genes across ovarian cancer cell lines and suggest ovarian cancer spheroids derived from HGS lines have consistent enrichment of SOX2, OCT4 and NANOG that correlate with traditional surface markers of TICs.

**Figure 3.**
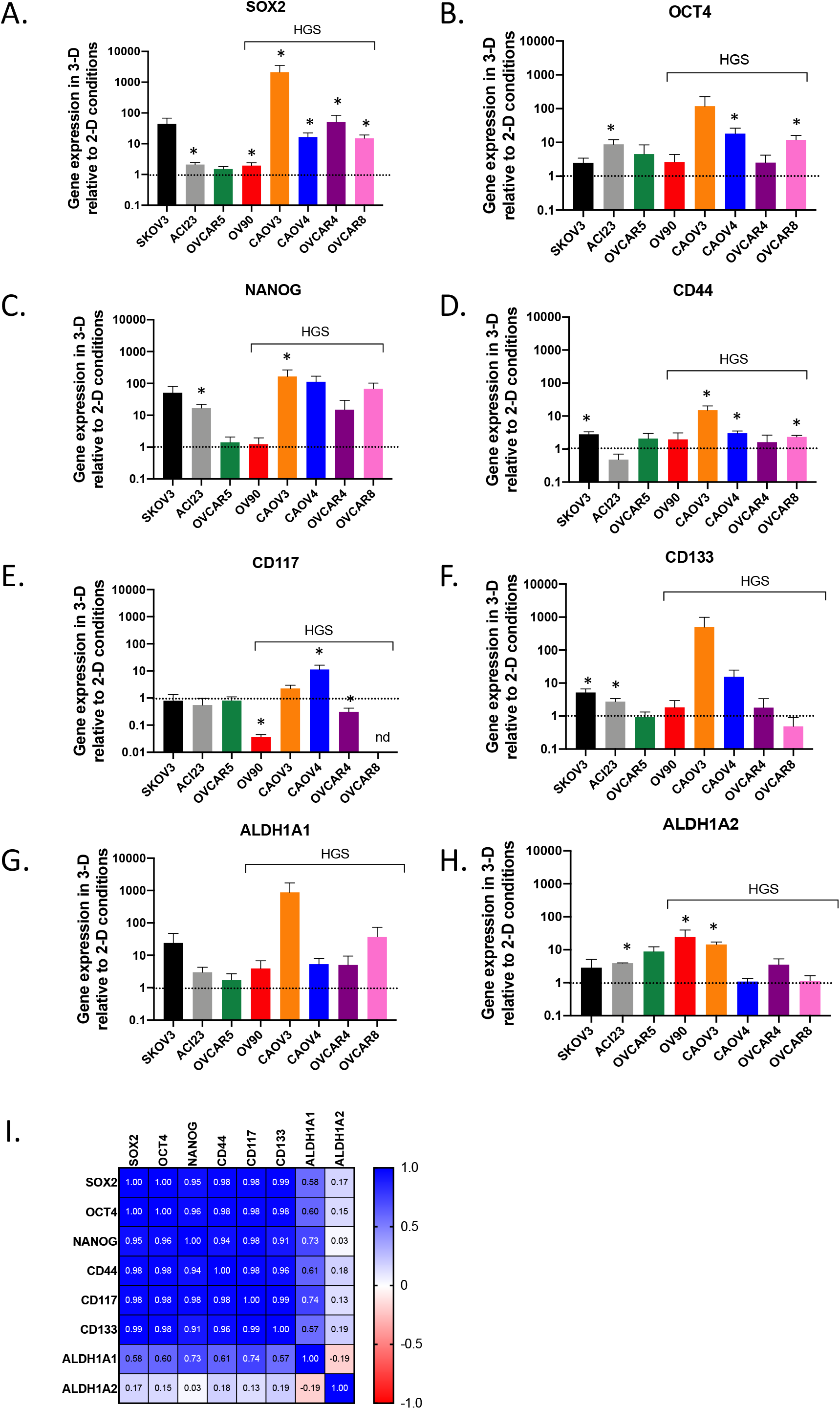
Gene Expression in ovarian cancer cells cultured in 3-D relative to 2-D conditions. Gene Expression data of cells grown in 3-D conditions compared to 2-D conditions for A) SOX2 B) OCT4 C) NANOG D) E) CD117 F) CD133 G) ALDH1A1 and H) ALDH1A2; I) Pearson correlation analysis of genes enriched in 3-D relative to 2-D conditions. Graphs represent mean and SEM. nd= not detected. Students T-test 3-D vs 2-D * p<0.05

Although TICs are enriched in 3-D culture conditions they remain a minority of the total population of cells [15]. We therefore wanted to establish expression levels of SOX2, OCT4, and NANOG in TICs isolated from 3-D cultures via FACS sorting based on traditional TIC markers. We chose to evaluate three cell lines from our panel: CAOV4 as a representative HGS line, ACI23 as an undefined serous line with exceptional growth in 3-D conditions, and OV90, a likely HGS line whose growth characteristics and SOX2, OCT4 and NANOG expression in 3-D are distinguishable from the other HGS lines in our panel. Each cell line was sorted for either high ALDH activity or high CD117, CD133 or CD44 expression (**Fig. 4A and Fig. S1A-E**). The percentage of CD44+ cells was significantly higher in OV90 cells relative to ACI23 and CAOV4 cells. The percentages CD117+ cells were less than 10% and not significantly different among all three cell lines. ACI23 cells had a significantly higher percentage of CD133+ cells relative to OV90 and CAOV4. Similar to CD44 expression, ALDH activity was highest in OV90 followed by CAOV4 and lowest in ACI23 cells. ALDH activity may be the result of ALDH1A1 or ALDH1A2 or a combination of both isoforms, as we have previously demonstrated [25]. These data again highlight the heterogeneity of ovarian TIC populations across cell lines and supplement previous findings demonstrating differential expression of TIC markers in OVCAR5, SKOV3, and A2780 cell lines [18, 47–49]. Quantification of SOX2, OCT4, and NANOG in FACS sorted marker positive populations relative to marker negative populations revealed that CD44+ cells did not have increased SOX2, OCT4, and NANOG, except in the ACI23 line, which had significantly elevated levels of OCT4 and NANOG (**Fig. 4B**). CD117+ cells derived from all three cell lines had enriched expression of SOX2, OCT4, and NANOG, although only ACI23 reached statistical significance (**Fig. 4C**). In contrast, CD133+ cells had no enrichment of SOX2, OCT4 or NANOG (**Fig. 4D**). The CAOV4 line had a negligible number of CD133+ cells and thus could not be analyzed. ALDH+ cells had no significant enrichment of SOX2, OCT4 or NANOG, except for ACI23 cells which had elevated SOX2 (**Fig. 4E**). These results indicate that although SOX2, OCT4, and NANOG are enriched in 3-D cultures, their expression is limited to a small population of CD117+ cells and/or cells expressing a combination of traditional markers, and highlights the need for more determinative TIC markers than those traditionally used.

**Figure 4.**
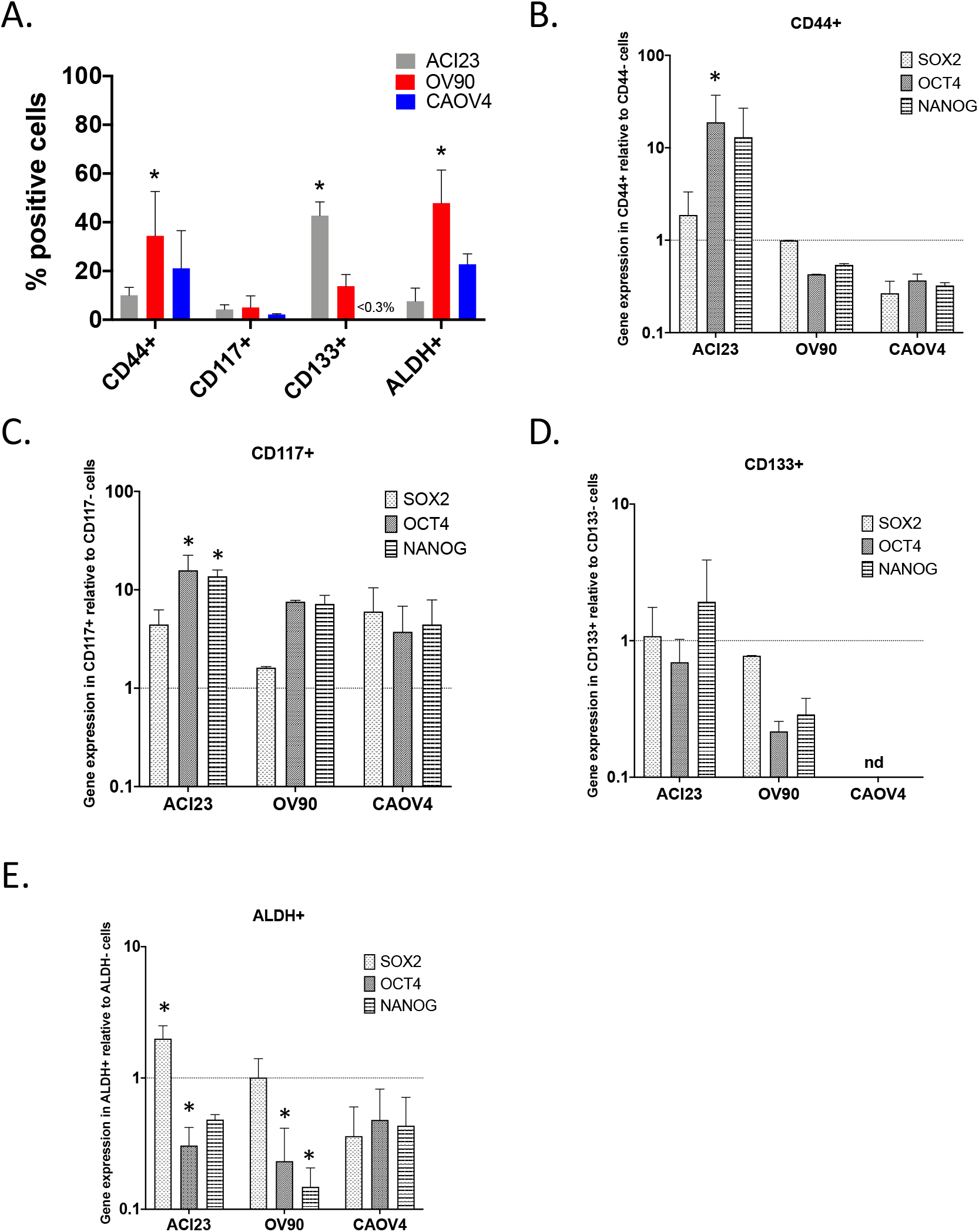
SOX2, OCT4 and NANOG in FACS sorted ovarian cancer cells. A) Percent positive cells retrieved from 3D cultured cells sorted using common stem cell markers (ALDH activity or CD133, CD44 or CD117 expression). B) Gene expression in ALDH+ cells relative to ALDH-cells. C) Gene expression in CD44+ cells relative to CD44-cells. D) Gene expression in CD133+ cells relative to CD133-cells. E) Gene expression in CD117+ cells relative to CD117-cells. Graphs represent mean and SEM relative to control *P<0.05 Two-way ANOVA. nd = not detected

Given the robust growth capacity of ACI23 cells and their consistent expression of SOX2, OCT4, and NANOG, we used this line to investigate in vivo growth in a limiting dilution assay and the corresponding endogenous expression of SOX2, OCT4 and NANOG. We found that mice that received 500K cells created tumors with 100% efficiency, whereas mice receiving 50K or 5K created tumors with 81.25% and 75%, efficiency, respectively, over a monitoring period of 120 days (**Fig. 5A**). Mice receiving 500 cells only developed tumors with 12.5% efficiency with only 2 tumors developing as late as day 46 and day 116 and were therefore not considered for further comparisons. Mice receiving 500K or 50K cells developed palpable tumors (time to tumor initiation) in 9-13 or 18-25 days, respectively (**Fig. 5B**). Mice receiving the more diluted concentration of 5K cells developed palpable tumors within a broader range of time (as early as 26 or as late as 45 days), suggesting a potential difference in tumor initiation efficiency within that group (**Fig. 5B**). Given the fundamental role of SOX2, OCT4, and NANOG in sustaining pluripotency and self-renewal in TICs, we were interested in determining the level of SOX2, OCT4, and NANOG in tumors that appeared early and thus had enhanced tumor initiation efficiency versus those that appeared late. Tumors from each dilution were ranked based on the number of days it took before palpable tumors appeared and were split into two equal groups of early and late appearing tumors. There was no significant difference in time to initiation in early or late appearing tumors in mice that received 500K or 50K cells; however, time to initiation was significantly longer in late appearing tumors in mice that received 5K cells (**Fig. 5B**).

**Figure 5.**
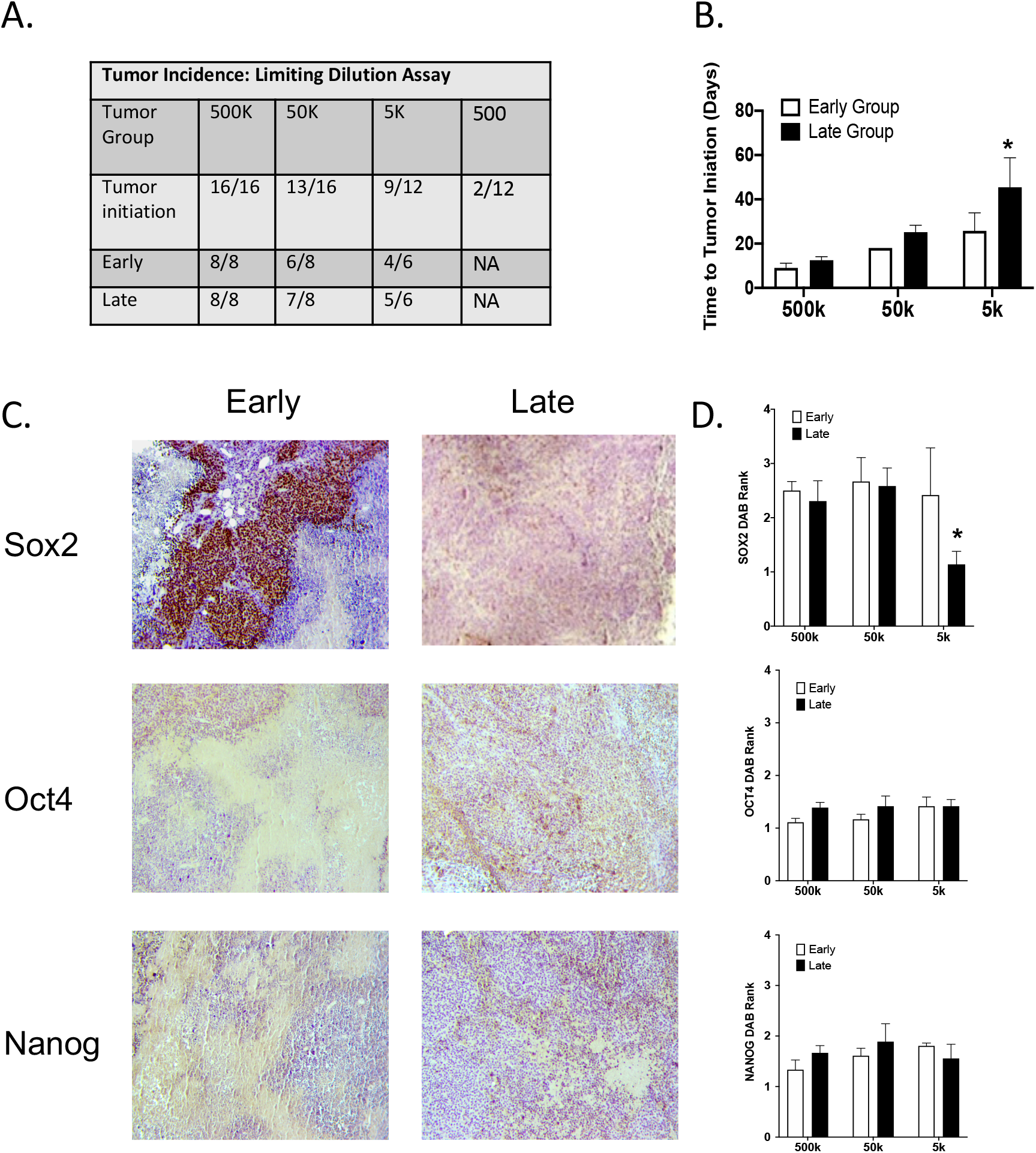
SOX2, OCT4 and NANOG in Early and Late Forming Tumors. A) Limiting dilution assay comparing early and late initiation of OV90 subcutaneous tumors. Appearance of palpable tumors (100mm^3^) were ranked and defined as Early (First half) or Late (Last half). B) Early and late tumor Initiation were significantly different when 5k or fewer cells were injected. C) Representative images of immunohistochemical staining of SOX2, OCT4, and NANOG in fixed tumor sections derived from early and late appearing tumors. D) SOX2, OCT4 and NANOG staining was quantified through ranking of intensity by three blinded investigators. Graphs represent mean and SEM *P<0.01. Two-way ANOVA, Bonferroni multiple comparisons testing.

Although tumors appeared early or late, all were resected after they reached an average volume of 879mm^3^. We then assessed SOX2, OCT4, and NANOG expression in early and late appearing tumors by immunohistochemical staining (**Fig. 5C**). SOX2 expression was not significantly different in early or late appearing tumors developed from 500K or 50K dilutions, however it was significantly increased in early appearing tumors developed from the 5K dilution (**Fig. 5D**). There was no significant difference in OCT4 or NANOG expression in early or late appearing tumors from any of the dilutions. These data suggest SOX2, relative to OCT4 or NANOG, is more strongly associated with tumor initiation efficiency, a fundamental feature of TICs, and may be a driver of tumor recurrence.

To begin to explore this possibility, we evaluated SOX2, OCT4 and NANOG expression in our panel of cell lines after chemotherapy treatment. We measured viability over 48 hours in 2-D and 3-D conditions across a range of carboplatin and paclitaxel concentrations and generated response curves (Fig. 6A-B). IC50 concentrations calculated for each cell line show that although they were variable across the cell lines in 2-D conditions, the HGS lines, relative to the other lines, had consistently higher IC50s in 3-D conditions for both carboplatin and paclitaxel. The exception is CAOV3 which was highly sensitive to chemotherapy and had the lowest IC50 among all the HGS lines. Using IC_30_ concentrations of carboplatin and paclitaxel we treated 2-D cultures for 72 hours and examined SOX2, OCT4 and NANOG expression (**Fig. 6D-F**). Relative to vehicle treated cultures all cell lines, except OVCAR5, OV90, and OVCAR4 showed enhanced expression of at least one of the genes, although SOX2 was the only gene significantly elevated (**Fig. 6D**). In order to more closely mimic clinical regimens, we extended these studies in 2-D cultured ACI23 cells that received 3 sequential treatments of IC_30_ carboplatin to evaluate SOX2, OCT4 and NANOG after multiple exposures (**Fig. 6G**). After two treatments with carboplatin, SOX2, OCT4, and NANOG all remain significantly elevated relative to pre-treatment levels; however, after 3 treatments of carboplatin only SOX2 remains significantly elevated, whereas OCT4 and NANOG are significantly decreased. Taken together, these data indicate a stronger role for SOX2 in chemotherapy resistance, a central feature of TICs that may contribute to cancer recurrence.

**Figure 6.**
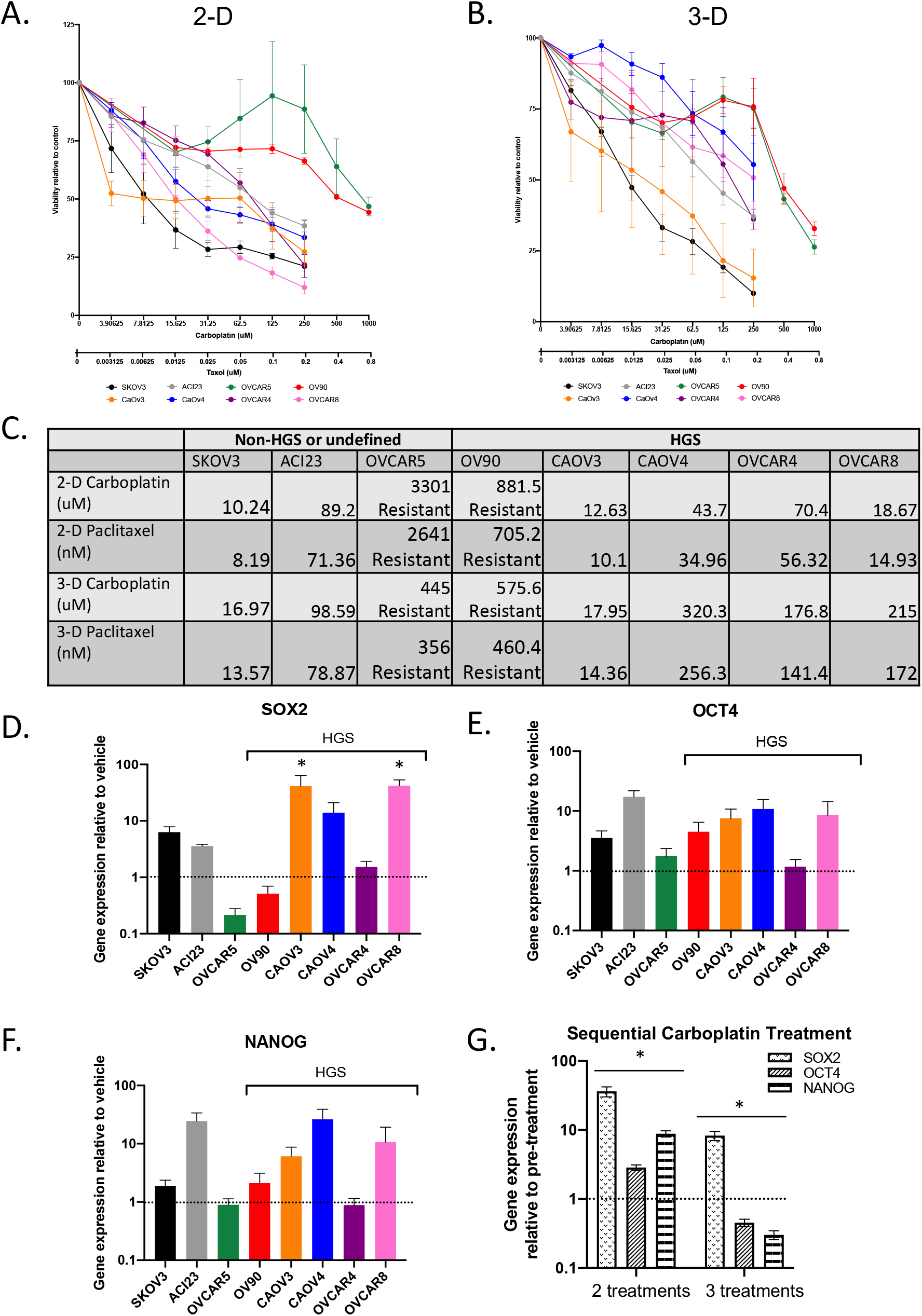
SOX2, OCT4, and NANOG in cell lines receiving chemotherapy. Viability growth assay of cells grown over 48 hours in combination (Carboplatin + Paclitaxel) treatment compared to vehicle control for A) 2-D conditions and B) 3-D conditions; C) IC50 calculations for Carboplatin or Paclitaxel in 2-D and 3-D conditions, calculated with Least Squares Fit; Gene Expression data of Combination (Carboplatin + Paclitaxel) treated relative to vehicle control D) SOX2 E) OCT4 F) NANOG; G) Gene Expression data of sequential carboplatin treatments relative to vehicle control. Graphs represent mean and SEM. Students T-test treated vs vehicle * p<0.05

To elucidate the clinical implications of our findings, we assessed the prognostic significance of SOX2, OCT4, and NANOG gene expression in TCGA ovarian cancer cases classified as recurrent (n=96) or disease-free (n=41) [34, 35, 50]. SOX2 expression was significantly higher in recurrent cases compared with disease-free cases (**Fig. 7A**); however, there was no significant difference in expression of OCT4 or NANOG in disease-free versus recurrent cases (Fig. **7B-C**). These data support our in vivo and in vitro findings indicating a stronger role for SOX2, relative to OCT4 and NANOG, in tumor initiation and drug resistance features of ovarian TICs. Thus, SOX2 may serve as an additional, and potentially more reliable marker of ovarian TICs to better distinguish ovarian cancer cells with high relapse potential.

**Figure 7.**
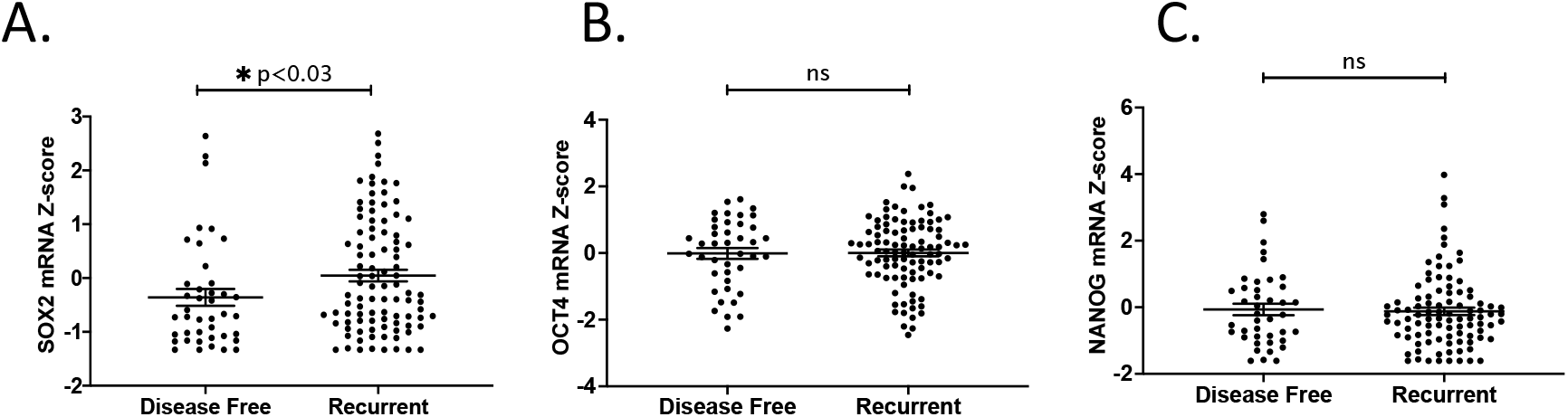
SOX2, OCT4 and NANOG in disease-free and recurrent patient tumors. mRNA Z-score and disease status of patients from HGSOC TCGA data set, Students T-test, for A) SOX2, B) OCT4, and C) NANOG.

## Discussion

In this study, we ultimately established SOX2 as a strong indicator of ovarian TICs. We found that across a variety of ovarian cancer cells, SOX2 expression correlates with growth in 3-D culture, spheroid formation capacity, chemoresistance, and expression of TIC surface markers. Our in vivo studies show that SOX2, relative to OCT4, and NANOG, correlates with early tumor onset. Finally, we used TCGA datasets to show that SOX2, but not OCT4 and NANOG expression, is significantly elevated in recurrent disease.

Our previous studies demonstrated that altered drug metabolism and oxidative stress pathways support growth of ovarian cancer spheroids. We now aimed to clarify the role of cell cycle pathway genes, including *SOX2*, that also support 3-D spheroids. *SOX2, OCT4*, and *NANOG* encode master embryonic transcription factors that are vital for quiescence, pluripotency, and long-term self-renewal [21, 51], properties characteristic of stem-like behavior. Expression of these genes has not been fully established in ovarian cancer TICs, so we took a comprehensive approach to characterize their expression in a panel of ovarian cancer cells cultured under different growth and treatment conditions. We also wanted to establish any correlations with traditional surface proteins and ALDH activity commonly used to isolate TICs. Given the heterogeneity of ovarian cancer cell lines and patient samples, these data would provide for more reliable identification of TICs with stem-like properties.

We found that all eight cell lines tested grew spheroids and had elevated SOX2, OCT4, and NANOG expression in 3-D conditions as compared to 2-D conditions. The five HGSOC lines had the greatest increase in SOX2, OCT4, and NANOG in 3-D, as well as the greatest ability to produce spheroids in 2-D conditions, as compared to the non-HGS or undefined lines. CAOV3 in particular had the highest enrichment of SOX2 in 3-D growth, in agreement with previous studies of CAOV3 spheroids [19]. SOX2 reached significance across all the confirmed HGSOC lines, while OCT4 and NANOG expression was variable. In our analysis of TIC marker gene expression across the cell lines, we found that, with the exception of CAOV3, most lines had less consistent enrichment of the genes encoding surface markers relative to their enrichment of SOX2, OCT4, and NANOG, although the HGS lines had the most enrichment. Interestingly, the *ALDH1A2* gene, unlike the other markers including *ALDH1A1*, was less enriched in the HGS but more enriched in the undefined or OV90 HGS line. Indeed, expression of this gene did not correlate with expression of any of the other genes investigated, highlighting the potential dependence of certain lines on different ALDH isozymes [52]. These data are in agreement with our previous work showing ALDH1A2 having higher expression and a stronger role in chemoresistance of ACI23 and OV90 ovarian TICs [25]. In marker sorted populations, only the CD117+ cells had enriched expression of SOX2, OCT4, and NANOG relative to its negative counterpart. Cells positive for CD44, CD133 or ALDH did not show consistent enrichment of SOX2, OCT4, and NANOG, despite its correlation with expression in 3-D conditions. Although these studies were only completed on three cell lines in our panel, the findings suggest SOX2, OCT4, and NANOG do not correlate with marker expression at the protein level or are only enriched in cells that express a combination of markers, such as CD133 together with ALDH, which is a commonly used marker combination for ovarian TICs [16].

In order to correlate SOX2, OCT4 and NANOG with long-term self-renewal properties in vivo, we evaluated their expression in a limiting dilution tumor model and found significant differences emerge between early and late tumors in the 5k dilution group relative to more concentrated dilutions. Tumors that appeared early had significantly higher expression of SOX2 relative to late appearing tumors. No differences in OCT4 or NANOG were observed for early or late tumors, suggesting a stronger role for SOX2 in this process. These data support other studies demonstrating the critical role of SOX2 in ovarian cancer progression [19, 24, 53]. Finally, relative to OCT4 and NANOG, we found that SOX2 remained elevated in chemotherapy resistant cells that survived multiple treatments of carboplatin in vitro. These findings suggest that SOX2 is a critical marker of chemoresistance that may indicate TICs capable of establishing relapsed disease. We investigated whether current clinical data in the TCGA database could corroborate this finding and we found that indeed, SOX2, but not OCT4 or NANOG, is significantly higher in tumors that were recurrent. These data are also in alignment with other studies showing that SOX2 is associated with worse overall survival of serous epithelial ovarian cancer [54–56]. Although additional studies are required to confirm SOX2 as a bona-fide TIC marker, our data confirms previous studies establishing SOX2 in spheroid formation and tumor initiation [19, 24] and indicates SOX2 is strongly correlated with a TIC phenotype and may serve as a more reliable marker of ovarian TICS with high relapse potential.

In addition to characterizing SOX2, OCT4, and NANOG in ovarian TICs, our work confirms the heterogeneity of ovarian cancer cells and TIC properties [14, 26–28]. Recent genomic analyses of ovarian cancer cell lines have demonstrated that some cell lines often used in research do not represent the genetic profile or ovarian tumor type with which they are often associated [29–31]. We found significant differences between the HGSOC lines and the non-HGSOC or undefined lines in viability, spheroid formation, chemoresistance, and gene expression in 3-D versus 2-D conditions. Additionally, the percentage of CD133+, CD44+, or ALDH+ cells ranged from <0.3% to 47.8% of the population. Our data show that CD117 may represent the most reliable marker for identifying cells with high SOX2, OCT4 and NANOG across cell lines, regardless of histological type. Moreover, CD117 is expressed on a minority of cells in all three lines tested. Due to their increased quiescent nature, we expect TIC populations to represent a small minority of the total population. CD117 is often used in combination with other markers, such as CD133 or CD44, to isolate ovarian TICs [48, 57]. Although this may enrich for a larger percentage of the cells, our data suggest that CD117 alone might be sufficient and potentially more reliable indicator of TICs with stemness properties associated with SOX2, OCT4 and NANOG. Our data is in agreement with a previous study of ovarian cancer xenografts showing that CD117+ cells were less than 2 percent of the total tumor cells and transplantation assays required 100 fold fewer CD117+ cells relative to CD117-cells [58]. Our data, combined with other published studies, strongly suggest CD117 may be a more reliable single marker for identifying ovarian TICs, and their expression of SOX2, OCT4, and NANOG may contribute to the quiescence and long-term self-renewal associated with these cells.

In conclusion, ovarian cancer has proven difficult to model due to its complex heterogeneity and poor clinical outcomes. The majority of patients are prone to therapy-resistant disease and tumor relapse within five years of diagnosis [1, 2]. Over a decade of work has identified TICs as a small subset of chemoresistant cells that may be responsible for relapse in many cancer types, but they remain a challenge to study due to their heterogeneity and potential lack of functional relevance. We sought to clarify whether SOX2, OCT4, and NANOG expression may improve classification of ovarian TICs with enhanced quiescence and long-term self-renewal. We found that elevated SOX2 expression best correlated with the TIC phenotype across cell lines, in tumor initiation studies in mice, and in TCGA studies for recurrence. Future efforts to study mechanisms of chemoresistance and tumor relapse in ovarian cancer will benefit from an improved method of TIC validation. A better understanding of TIC biology will lead to more effective therapeutic strategies for preventing relapse and prolonging remission for ovarian cancer patients.

## Acknowledgments

We thank Ruby Tandon of the SDSU FACS core facility for her excellent assistance with flow cytometry experiments. This work was funded by the National Cancer Institute at the NIH under award number R00CA204727 and the National Institute on Minority Health and Health Disparities award number U54MD012397. The authors declare no conflict of interest.

## Supplemental Figure 1

**Figure S1.**
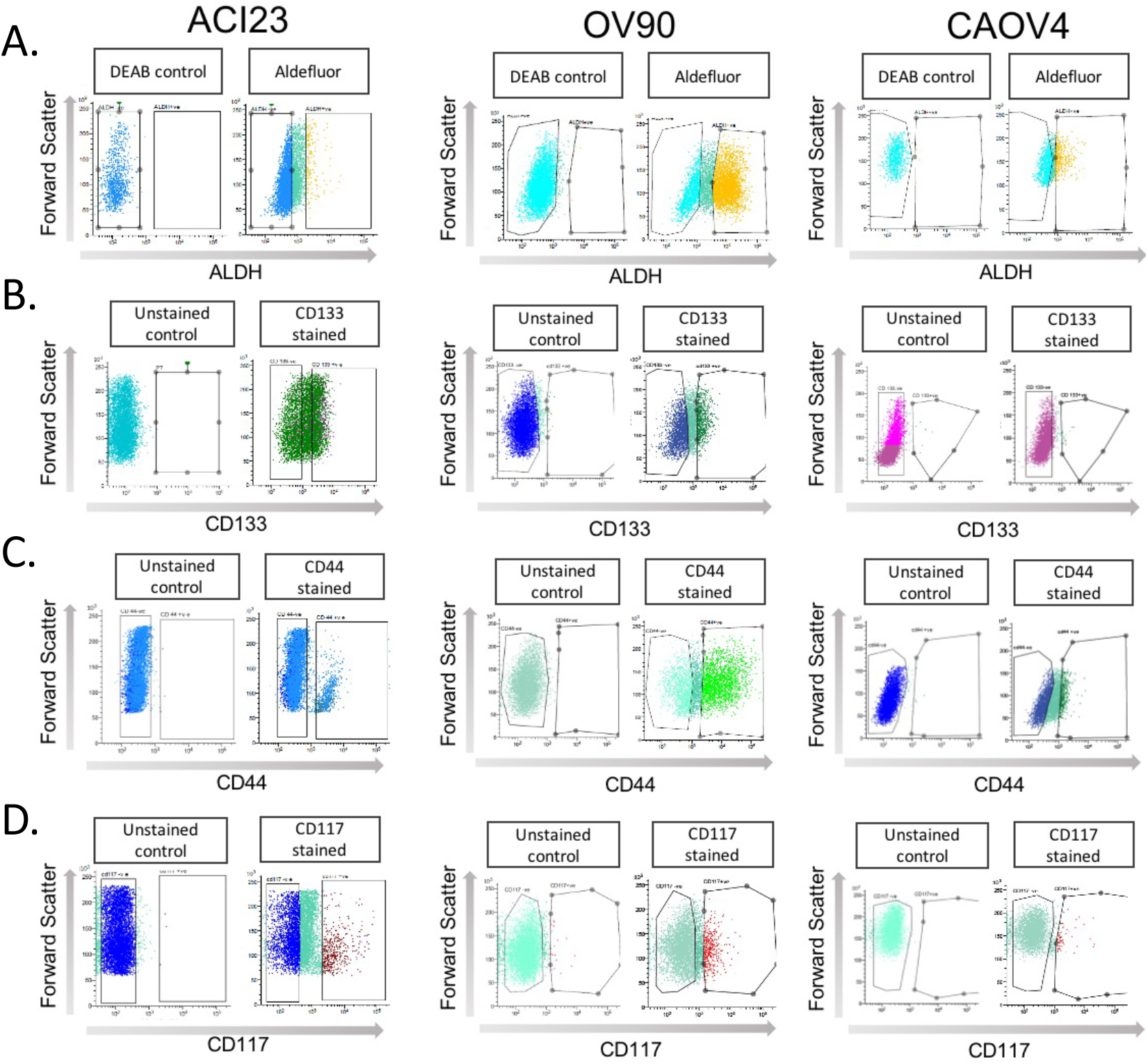
Gating strategy for FACS Experiments. Representative gates for sorting of ACI23, OV90, and CAOV4 ovarian cancer cells with high ALDH activity (A), CD133 expression (B), CD44 expression (C), or CD117 expression (D).

## Notes

### Competing Interest Statement

The authors have declared no competing interest.

